# Separable neural mechanisms support intentional forgetting and thought substitution

**DOI:** 10.1101/2020.10.29.360511

**Authors:** Ryan J. Hubbard, Lili Sahakyan

**Author notes:** Corresponding author: Ryan J. Hubbard.

## Abstract

Psychological and neuroscientific experiments have established that people can intentionally forget information via different strategies: direct suppression and thought substitution. However, few studies have directly compared the effectiveness of these strategies in forgetting specific items, and it remains an open question if the neural mechanisms supporting these strategies differ. Here, we developed a novel item-method directed forgetting paradigm with Remember, Forget, and Imagine cues, and recorded EEG to directly compare these strategies. Behaviorally, Forget and Imagine cues produced similar forgetting compared to Remember cues, but through separable neural processes; Forget cues elicited frontal oscillatory power changes that were predictive of future forgetting, whereas item-cue representational similarity was predictive of later accuracy for Imagine cues. These results suggest that both strategies can lead to intentional forgetting, but directed forgetting may rely on frontally-mediated suppression, while thought substitution may lead to contextual shifting, impairing successful retrieval.

## Introduction

For many people and situations, forgetting is considered a negative experience, and is something that one rarely does on purpose. However, in some instances forgetting is as critical as remembering; we must often forget outdated information, such as where we parked our car yesterday, or more negative or traumatic events that are painful to recall. More often than we realize, forgetting can be the goal and serve a positive function, and thus it is important to understand the mechanisms that produce successful intentional forgetting. A variety of laboratory paradigms have been developed to study the mechanisms of intentional forgetting, including the directed forgetting (DF) procedure (Bjork, LaBerge, & LeGrand, 1978) and the think-no-think (TNT) procedure (Anderson & Green, 2001). These procedures involve instructing participants to forget or to not think of some previously presented information. Decades of research have shown that participants have worse memory for items they are told to forget or not think about (for reviews, see Anderson & Hanslmayr, 2014; Sahakyan, in press; Sahakyan, Delaney, Foster et al., 2013), suggesting that we can voluntarily control our memory, and that we can impair access to the unwanted information. However, the question of what specific mechanisms give rise to this impaired memory for unwanted information remains a much debated topic in the literature. The goals of the current investigation were not only to assess the effectiveness of thought substitution as a strategy for intentional forgetting of individual items, but also to directly compare the neural mechanisms of directed forgetting and thought substitution to better understand the mechanisms of intentional forgetting.

Multiple studies from our lab have investigated what people do when they are given a cue to forget information in list-method DF studies (for a review, see Sahakyan et al., 2013; Sahakyan & Foster, 2016). Although a variety of controlled strategies are reported by the participants, *thought substitution* (also known in the literature as diversionary thought or mental context-change) is one of the most effective strategies for impairing memory recall, akin to what is typically observed with a more standard Forget cue (Delaney, Sahakyan, Kelley et al., 2010; Foster & Sahakyan, 2011; Sahakyan & Kelley, 2002). For instance, when participants were instructed to imagine being invisible (Sahakyan & Kelley, 2002), think about their childhood home (Sahakyan & Delaney, 2003), daydream about a vacation (Delaney et al., 2010), or wipe a computer monitor (Mulji & Bodner, 2010) between the two lists, DF-like effects were observed, even without any explicit instruction to forget. Thought substitution instructions not only produced impaired memory for the first list of items, but also these effects were found irrespective of encoding strategies (Sahakyan & Delaney, 2003), as well as across individual differences in working memory capacity (Aslan, Zellner, & Bäuml, 2010; Delaney & Sahakyan, 2007), across age differences including in children and older adults (Aslan & Bäuml, 2008; Sahakyan, Delaney, & Goodmon, 2008), across serial position effects in memory (Sahakyan & Foster, 2009), and across nuanced measures of retrieval dynamics (Unsworth, Spillers, & Brewer, 2012). These results suggest that thought substitution can be a powerful strategy for intentional forgetting of information. At the same time, recent evidence seems to suggest that the mechanisms underlying directed forgetting and thought substitution may not be the same, as memory impairments from directed forgetting persist across delay, whereas thought substitution effects dissipate with delay (Abel & Bäuml, 2017; Hupbach, 2018). Additionally, thought substitution strategies have been shown to be effective in list-method paradigms, but no study to date has investigated the utility of thought substitution in an item-method paradigm.

Studies measuring only behavioral responses may identify similarities or differences between directed forgetting and thought substitution in overall accuracy or reaction time, but may lack the sensitivity to pinpoint the specific differences in neural mechanisms engaged by these strategies at the level of individual trials, particularly at the time of processing the forget or substitution cue. However, utilizing non-invasive brain measurements while participants engage in intentional forgetting may elucidate the different mechanisms engaged by different strategies.

Previous studies have examined whether intentional forgetting through directed forgetting differs in neural processing from passive forgetting. For instance, fMRI studies indicate that item-method DF engages an active process that suppresses ongoing encoding, often through activity in dorsolateral prefrontal cortex (DLPFC) and the hippocampus (Anderson & Hulbert, 2020; Butler & James, 2010; Depue, Curran & Banich, 2007; Reber, Siwiec, Gitleman et al., 2002; Nowicka, Marchewka, Jednorog et al., 2011; Rauchs, Feyers, Landeau et al., 2011; Rizio & Dennis, 2013; Wylie, Foxe & Taylor, 2008). EEG studies also suggest that successful intentional forgetting involves different neural mechanisms than passive, incidental forgetting, with Remember cues eliciting rapid ERP differences and Forget cues eliciting later frontal activity (Gallant & Dyson, 2016; Paller, 1990; Paz-Caballero, Menor & Jiménez, 2004; Ullsperger, Mecklinger & Müller, 2000; Van Hooff & Ford, 2011; Van Hooff, Whitaker & Ford, 2009). One simultaneous EEG-fMRI study found that Forget cues led to increases in DLPFC activity along with decreased neural phase synchrony, and transcranial magnetic stimulation to the DLPFC reduced successful intentional forgetting (Hanslmayr, Volberg, Wimber et al., 2012). Additionally, intracranial EEG studies have provided more direct evidence that the hippocampus (Ludowig, Möller, Bien et al., 2010), as well as the DLPFC (Oehrn, Fell, Baumann et al., 2018) are involved in voluntary forgetting of information. In sum, neuroscientific studies of directed forgetting generally support the notion that intentional forgetting is an active process requiring cognitive resources.

While the neural mechanisms of directed forgetting have been well-studied, only a handful of cognitive neuroscientific studies have compared directed forgetting and thought substitution. EEG studies comparing these strategies in a TNT paradigm (Bergström, de Fockert & Richardson-Klavehn, 2009) and in a list-method DF paradigm (Bäuml, Hanslmayr, Pastötter et al., 2008; Pastötter, Bäuml & Hanslmayr, 2008) suggest that directed forgetting and thought substitution elicit different and opposing oscillatory power changes, with directed forgetting leading to alpha band increases and thought substitution eliciting alpha band decreases, and only directed forgetting leads to reduction of recollection-related ERPs. Additionally, fMRI evidence (Benoit & Anderson, 2012; Kim, Smolker, Smith et al., in press) suggests that separable neural networks support these strategies. Namely, prefrontal inhibition of hippocampal processing may support directed forgetting, whereas thought substitution involves recruitment of a left prefrontal cortical circuit. However, these studies have generally utilized list-method DF and TNT paradigms, and it remains unclear if thought substitution is an effective in-the-moment strategy for forgetting information, and whether separable neural mechanisms support these strategies at shorter timescales.

To better understand how these strategies differ in producing forgetting, we developed a novel item-method DF paradigm which included a thought substitution condition. Participants studied items to remember for a later memory test, and were told to either Remember the item, Forget the item, or Imagine a memory from their own past. This Imagine instruction has been shown to be effective in producing forgetting in list-method studies (Sahakyan & Kelley, 2002), but has not been used in an item-method study before. Additionally, we recorded EEG while participants studied the items and performed these instructions in order to compare the neural mechanisms supporting directed forgetting and thought substitution. We employed multiple analyses in order to examine these differences at different levels of neural processing. Namely, we analyzed ERPs to compare changes in evoked potentials between various conditions of the experiment. We also performed time-frequency analyses in order to examine changes in oscillatory power, and employed representational similarity analysis (RSA), which allows for measurement of the neural pattern similarity of different item representations (Kriegeskorte, Mur & Bandettini, 2008). This latter type of analyses was recently used to examine item-cue similarity in an item-method DF experiment (Fellner, Waldhauser & Axmacher, 2020) in order to understand the fate of item representations during cue processing. Finally, we used multi-level models to predict behavioral accuracy from these different measurements of neural processing in order to establish brain-behavior relationships.

## Materials and Methods

### Participants

44 right-handed individuals participated in the experiment in exchange for payment (5$ per half hour, generally 15$ for the whole experiment). This sample size was chosen based on previous neuroscientific studies of directed forgetting (e.g., Schindler & Kissler, 2018). Eight participants were dropped from the analysis due to noisy data (large amounts of movement artifacts, technical issues with electrodes) or poor memory performance (2 participants were very tired during memory testing and performed at chance level), resulting in a total of 36 participants in the final analysis. All participants reported normal or corrected vision and had no history of any neurological or psychiatric disorder. Mean age was 21 years (range 18-30 years), and 24 of the participants were female. The study was approved by the Institutional Review Board of UIUC, and all participants provided written informed consent and were debriefed following participation.

### Materials

The stimuli used in the experiment were 210 nouns retrieved from the MRC Psycholinguistic Database (Coltheart, 1981). The words were medium frequency (Kucera & Francis mean word frequency of M= 43, SD=18) and were 4-6 letters in length. The assignment of each word as either an old or new word, as well as to each of the three memory instruction conditions, was randomized for each participant.

Picture images were used to designate the three memory instructions – Remember, Forget, and Imagine. The pictures were downloaded from emojipedia.org, and are displayed in Figure 1. Remember cues were represented by a green check mark, while Forget cues were depicted by a red X. For the Imagine cues, pictures of a house, a utensil set, and an airplane were used.

**Figure 1.**
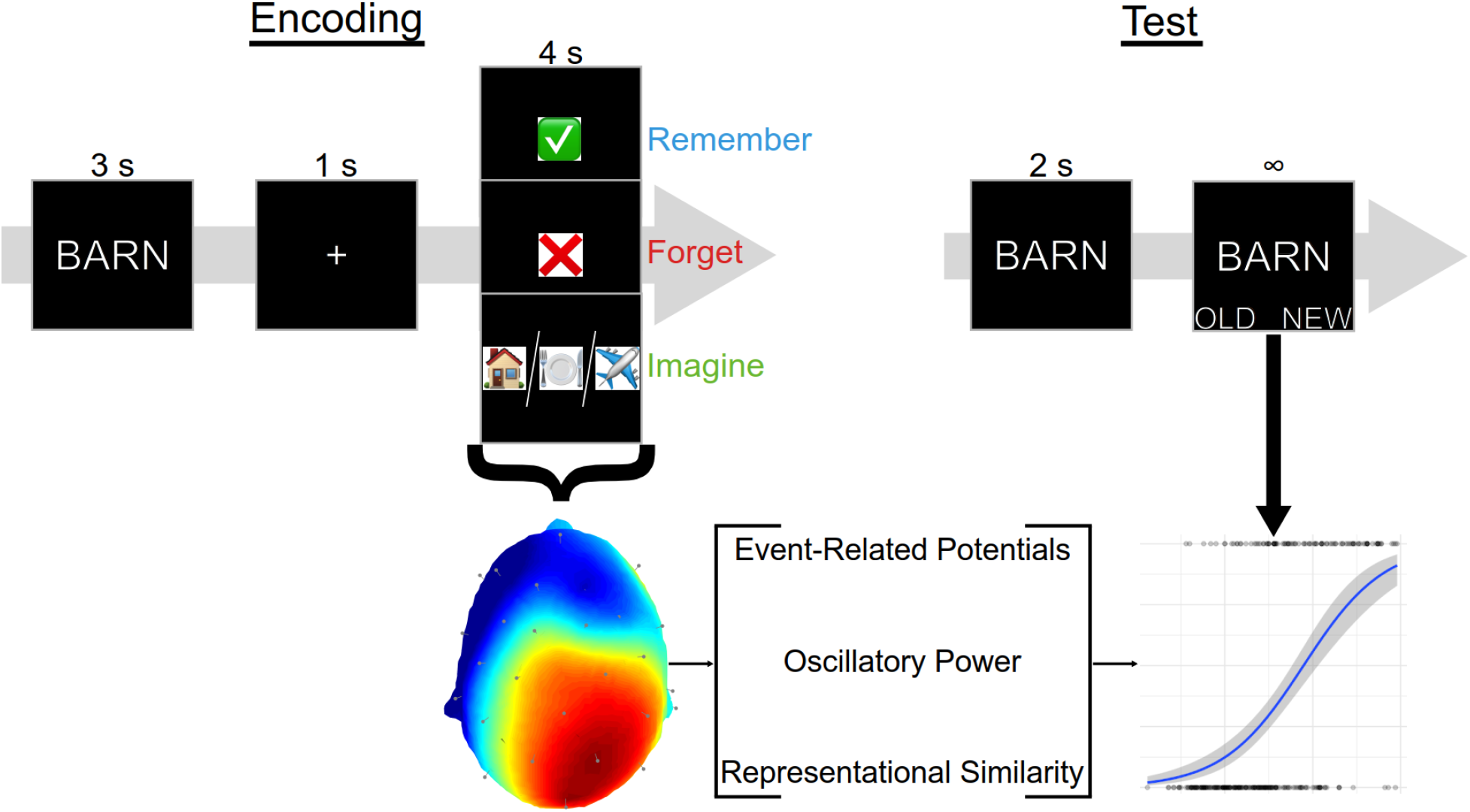
Outline of the experimental paradigm and analytical strategy. Participants studied words during encoding and were given either a Remember, Forget, or Imagine cue following each item. Following encoding, participants were given a recognition memory test on the words they had studied, as well as new words. EEG was recorded during the experiment, and differences between cues were assessed with multiple analyses. These neural differences were extracted at the trial level and submitted to mixed logit regression models predicting behavioral accuracy on the recognition test.

### Procedure

After informed consent and EEG setup, participants were comfortably seated approximately 100 cm from a monitor in a quiet room, where they received instructions for the DF task. Participants were told that they would view words to remember for later, but that only some words would be tested during the final memory test. They were told that if the Forget cue followed a word, it would mean that the word would not be tested and they should attempt to forget it. If the Remember cue followed a word, it meant the word will be tested and should be maintained in mind. Finally, participants were told that sometimes they might see the Imagine cue after a word, and that the rationale for such cues was to examine how attention shifts and mind wandering affect memory. Importantly, participants were told the words prior to Imagine cues would still appear on the memory test. After these instructions, participants were presented with each of the three separate Imagine prompts, and were given 60 s to visualize and verbally describe a clear mental image that related to the cue. The Imagine cues were selected from comparable DF studies that compared Forget condition to thought substitution/ mental context-change condition in list-method DF (e.g., Sahakyan & Kelley, 2002; Delaney et al., 2010). For the House cue, participants were told:

> *“I want you to imagine you are in your childhood home. Imagine you are entering through the front door, and visualize the house as you travel from room to room, including details about the furniture and their location. Please verbally describe the mental image you are seeing.”*

For the Cafeteria cue, participants received the following prompt:

> *“I want you to mentally travel back in time to high school. Imagine it’s time for lunch and you’re walking into the cafeteria, or whatever space you would generally eat at this time. Think of the people you are sitting with, the food you are eating, and the smells, sounds, and layout of the room.”*

Lastly, for the Vacation cue, participants were given the following prompt:

> *“I want you to think back to a vacation you took, or perhaps a trip for a class, and picture the things you saw and activities you participated in. Think of where you went, how you felt, who you were with, and the experiences you had on your trip. Please verbally describe the mental image now.”*

Participants were given practice trials with Imagine prompts to ensure familiarity with the task and to make it easier for them to shift into these mental contexts during the actual DF task. Three different imagine prompts (as opposed to a single prompt) were used to increase the chances of engaging in different mental contexts shifts throughout the experiment rather than repeatedly re-visiting the same mental context. All of the participants in the study were able to provide vivid images and details in response to the imagination prompts.

The participants were then instructed that they would be presented with a word to study, followed by an image that would tell them what to do next. If they saw the green check mark, they should try to remember the word, as their memory for it would be tested later. If they saw the red X, they should try to forget the word, and their memory for that word would not be tested later. Lastly, if they saw one of the three Imagine cues, they should create the mental image that they created before, and focus on creating that visual imagery, not on the word they just saw. Once the participant confirmed the instructions were clear to them, the encoding phase of the experiment began.

The outline of the procedure is displayed in Figure 1. On each trial of the encoding phase, a study word was presented centrally for 3 seconds, followed by a 1 second fixation, and then a 4 second cue presentation. The order of cue presentation was pseudo-randomized, such that trains of the 3 cues in a random order (R/F/I) were presented sequentially (e.g. I/F/R, R/I/F, etc.). Participants viewed 126 words and cues in total, with 42 in each cue condition, and 14 in each of the three Imagine conditions. Participants were instructed to focus on the words and perform the task instructed by the cue, and were not given any particular instructions on how to encode the words or any response to make on the keyboard. To reduce fatigue, participants were given 6 breaks lasting 15 seconds each throughout the encoding phase.

Following the encoding phase, the participants performed a recognition task, in which they indicated whether presented words were old or new. They were informed that their memory for the Forget cue items would in fact be tested, and they should respond “old” even if they recall that the word was originally a Forget cue item (i.e. the Forget instructions were canceled at the time of test). Participants were presented with all 126 study words, intermixed with 84 new words, for a total of 210 words in a random order. Each word was presented centrally for 2 seconds, after which a prompt to make an Old/New response appeared. Testing was self-paced. Participants responded whether the word was old or new with either a right hand or left hand response on the keyboard. The assignment of hand to response was counterbalanced across participants to avoid any potential confounds with neural responses Note that prior to the DF task, participants initially performed a stop signal task. The purpose of the stop signal task was to examine potential similarities in neural signals of stopping motor actions and inhibiting words in a DF task (Castiglione, Wagner, Anderson et al., 2019). The results of the stop signal task and their relationship to DF is not the focus of this paper and was not analyzed here.

### Behavioral Analysis

Recognition memory performance was analyzed with mixed effect logistic regression models. These models predicted whether participants made a correct or incorrect recognition response on trial-level behavioral data. Models were fit by maximum likelihood using the lme4 package in R (Bates, Mächler, Bolker et al., 2015), and Wald’s z-scores were computed for each coefficient to test for significance of fixed effects. Random factors included intercepts for items and slopes and intercepts for participants for the fixed effect of condition. Correlations between random factors were not calculated to ease convergence of the models. To test differences in recognition accuracy by cue condition, cue condition was included as a fixed effect in the model.

### EEG Recording and Pre-Processing

EEG data were recorded from 26 Ag/AgCl electrodes embedded into a flexible elastic cap and distributed over the scalp in an equidistant arrangement; see icon in Figure 3. Additional facial electrodes were attached for monitoring of electro-oculogram (EOG) artifacts, including one adjacent to the outer canthus of each eye and one below the lower eyelid of the left eye. Electrode impedances were kept below 10 kΩ. Signals were amplified by a BrainVision amplifier with a 16-bit A/D converter, an input impedance of 10 MΩ, an online bandpass filter of 0.016–100 Hz, and a sampling rate of 1 kHz. The left mastoid electrode was used as a reference for on-line recording; offline, the average of the left and right mastoid electrodes was used as a reference.

Following data collection and offline re-referencing, each raw EEG time series was passed through a 0.2–40 Hz Butterworth filter with a 36 dB/oct roll-off. Filter parameters were chosen a priori to remove low frequency drifts without causing artifacts in ERP analyses (Tanner, Morgan-Short, & Luck, 2015), as well as to remove high frequency noise but still allow for examination of beta band activity in time-frequency analyses. The time series was then segmented into epochs ranging from −700 to 2,500 ms relative to the onset of each encoding item and memory cue. While a time window of 2 seconds following each stimulus (either after a word or after a memory cue) was examined, significant effects were only observed within 1 second following each stimulus, and thus only results in the 1 second window are reported. Epoched data were then submitted to AMICA, an ICA algorithm that decomposes the signal into independent components (Palmer, Kreutz-Delgado, & Makeig, 2012). Each component timecourse was correlated with a bipolar vertical EOG channel (the lower eye channel – the channel above the left eye), as well as a bipolar horizontal EOG channel (the subtraction of the two outer canthus channels). Components with high correlations and topographies indicative of eye-related activity were removed, and the data was reconstructed from the remaining components. Lastly, the EOG-cleaned data was scanned for large voltage deflections (>90 μV), and manually scanned by eye, to remove any epochs with remaining artifacts. Overall, data quality was high, and few trials were removed (an average of 1.2% across subjects).

### ERP Analysis

ERP analyses were focused on changes in scalp voltage in response to stimuli. Prior to averaging, trials were baseline corrected with a z-score baseline procedure that reduces potential biases from standard baseline correction procedures (Ciuparu & Mureşan, 2016). The time series from −200 to −1 ms pre-stimulus was extracted from each trial and concatenated. Trials that were identified as artifacts were left out of the baseline. Each trial was then z-scored by the average and standard deviation of the concatenated baseline. Separate baseline corrections were performed for items and memory cues. Following baseline correction, trials were averaged to create ERPs.

Given the novelty of the experimental design, we chose to not select specific channel clusters or time-windows for statistical analysis of ERP components, but instead submitted the ERPs to time-constrained cluster-based permutation tests to identify significant differences between conditions (Maris & Oostenveld, 2007). Permutation tests with restricted time-windows increases statistical power while maintaining Type I error rate (Fields & Kuperberg, 2020). In these tests, t-tests were calculated at each time-point and channel, and significant t-values that were adjacent in space and time were clustered together. Clusters were characterized by taking the sum of t-values within the adjacent points. These observed clusters were compared to a permutation distribution, generated by shuffling the condition labels of the data, finding clusters, and summing the t-values of the clusters 2,000 times. Distributions of the most extreme cluster sums were created for comparison to the observed cluster sums. Reported p-values represent the percentile ranking of the observed clusters compared to the permutation distribution.

### Time-Frequency Analysis

Time-frequency analyses were focused on changes in oscillatory power in response to stimuli. Time-frequency decomposition was performed using FieldTrip functions (Oostenveld, Fries, Maris et al., 2011). EEG epochs were convolved with Morlet wavelets that varied in width (number of cycles) to improve temporal and frequency precisions. The width started at 3 and increased linearly to a width of 7 across a frequency range of 3 to 30 Hz, resulting in time-frequency bins of 20 ms and 0.5 Hz, respectively. Similar to the ERP analysis, the time-frequency data was then baseline corrected with the concatenated z-score procedure. A baseline period of −400 to −200 ms was extracted from each trial and at each frequency, and the concatenation of this baseline data was used to z-score the trial data. The trials were then averaged together. All statistical analyses of time-frequency data were carried out with non-parametric cluster-based permutation tests (Maris & Oostenveld, 2007).

### Representational Similarity Analysis

We used representational similarity analysis (RSA) to examine neural pattern similarity between encoding item activity and its associated cue. For each trial during the encoding phase, the vector of channel activity at each time-point following the encoding stimulus was correlated with the vector of channel activity of every time-point after the following memory cue, producing a matrix of correlations with the matching time-points on the diagonal. Here, 1000 ms of the encoding stimulus activity was correlated with 1000 ms of the memory cue activity. However, the data was downsampled to 250 Hz prior to RSA to reduce computational intensity and increase statistical power. The resultant matrices were averaged across trials for each cue condition, and the differences in neural similarity were statistically tested by with cluster-based permutation statistics.

### Behavioral Prediction

Memory researchers often investigate brain-behavior relationships in episodic memory studies by examining “Dm effects”, or separating and averaging activity at encoding based on later memory success (Paller & Wagner, 2002). However, these analyses may not specifically identify activity that is “predictive” of memory success, and predictive modeling approaches may be more effective (Chakravarty, Chen & Caplan, in press). Thus, we investigated how predictive the observed brain signals that differentiated conditions were of memory accuracy using mixed-effects logistic regression models (Jaeger, 2008). On *each* trial for each participant, mean signals were extracted from the significant clusters identified in the ERP, time-frequency, and RSA analyses. These signals were then entered into mixed-effect logit models predicting accuracy of individual trials (correct vs. incorrect response), with random intercepts for participants and items, as well as random slopes for participants for each of the brain signals. Statistics of the model fixed effects, including estimates, standard errors, and significance, are reported.

## Results

### Behavior

Participants’ recognition memory performance for the four types of test items (Remember, Forget, Imagine, and New) are plotted in Figure 2A. Participants performed above chance, and were able to successfully correctly reject New items. The mixed logit model predicting memory accuracy revealed significant differences between cue conditions; namely, participants had higher memory accuracy for Remember cue items than Forget (*β* = 1.21, *z* = 8.65, *p* < 0.01) as well as Imagine (*β* = 0.92, *z* = 5.76, *p* < 0.01) cue items. Additionally, accuracy for Imagine cue items was significantly greater than for Forget cue items (*β* = 0.29, *z* = 3.24, *p* < 0.01). Thus, we observed a directed forgetting effect for both Forget and Imagine items, but the magnitude of the effect was greater for Forget cue items.

**Figure 2.**
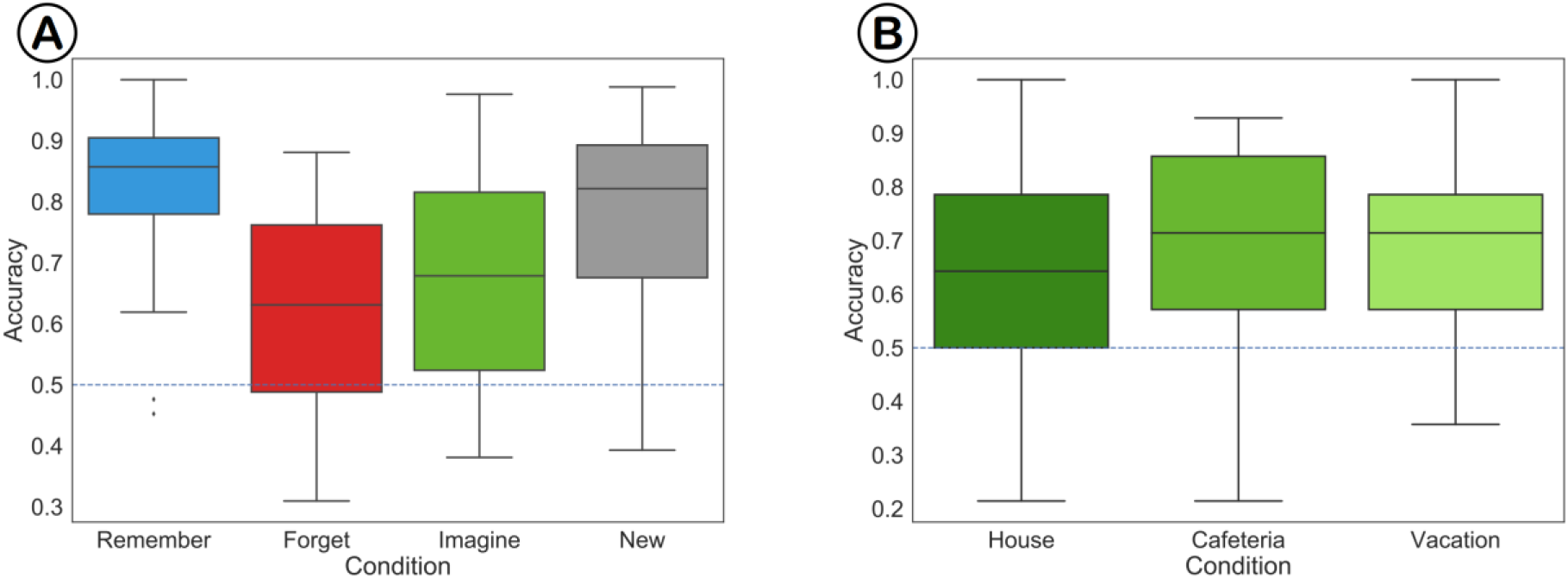
Recognition memory performance. Accuracy is plotted on the Y axis, and a line is drawn at chance performance (0.5 accuracy). A) Performance across the different item types. B) Performance across the different Imagine cue conditions.

To assess if performance in the Imagine cue condition may have been driven by one of the Imagine prompts over the others due to one of the prompts being potentially more likely to promote forgetting (e.g., House, Cafeteria, or Vacation), we examined accuracy across three sub-conditions. Accuracy for the three Imagine prompts was statistically analyzed with a mixed logit model using Imagine cues as a fixed effect (the results are plotted in Figure 2B). There were no significant differences between conditions (House vs. Cafeteria, *β* = 0.22, *z* = 1.31, *p* = 0.19; House vs. Vacation, *β* = 0.07, *z* = 0.44, *p* = 0.66; Cafeteria vs. Vacation, *β* = 0.15, *z* = 0.93, *p* = 0.35). Given that there were no significant differences between Imagine sub-conditions, in subsequent electrophysiological analyses we collapsed across Imagine sub-conditions.

### ERPs

ERPs to memory cues at central and frontal channels are plotted in Figure 3. Cluster-based permutation tests found differences in amplitudes between the three conditions of interest.

**Figure 3.**
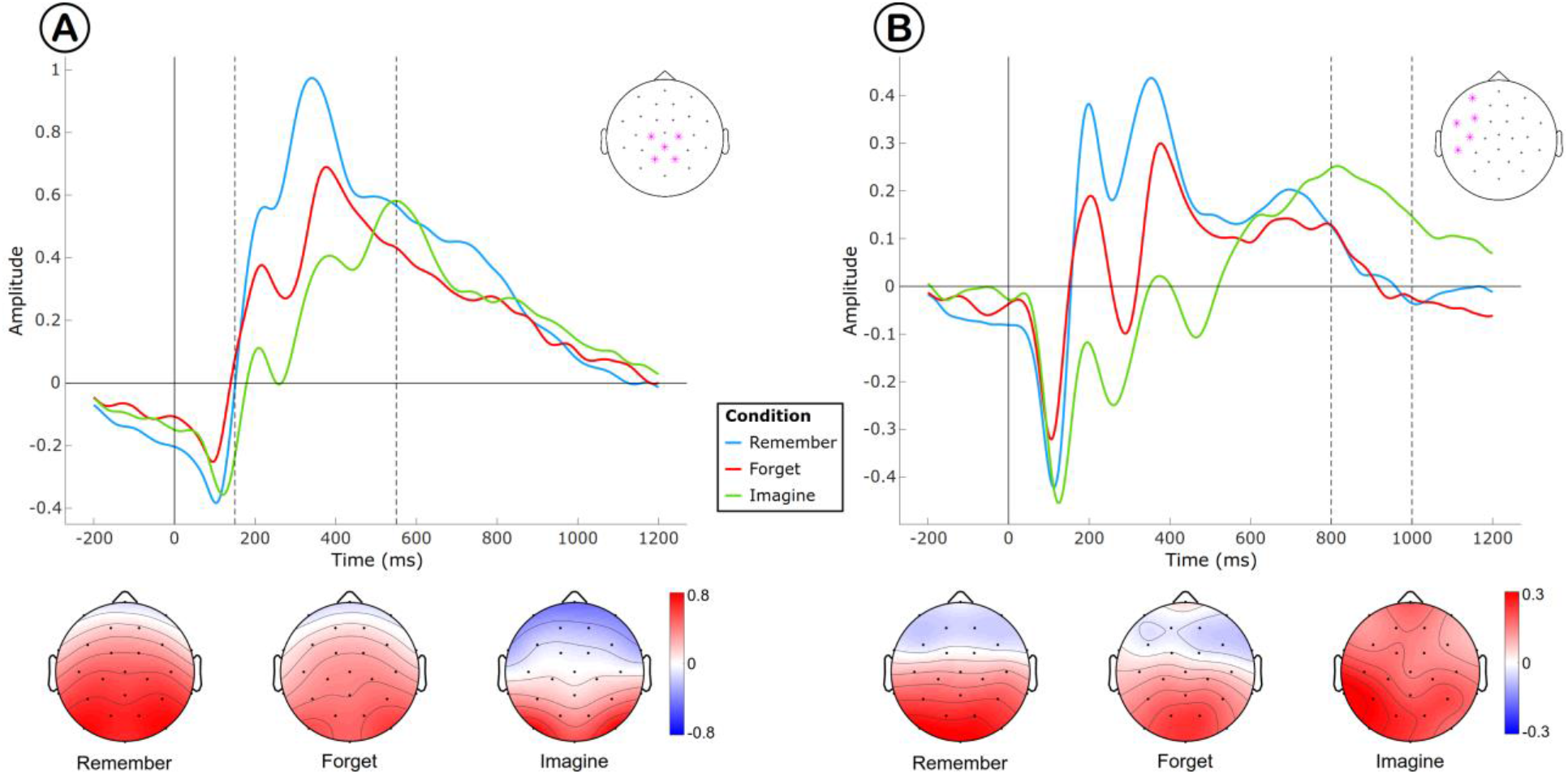
ERP results. ERPs time-locked to memory cues are presented at two channel clusters, a central cluster (A) and a left frontal cluster (B), where significant differences between conditions were found. Topographies of average scalp activity are shown below the ERP plots, and the time window for averaging is between the dotted lines of the ERP plot: 150-550 ms (A), and 800-1000 ms (B).

In the comparison of Remember vs. Forget cues, the cluster-based test revealed a significant difference in activity between 150 and 850 ms most pronounced over central and posterior channels, with Remember cues eliciting greater amplitude than Forget cues (*p* < 0.01). A second cluster was also found with a frontal distribution earlier in time, between 100 and 160 ms, with Forget cues eliciting higher amplitudes than Remember cues (*p* = 0.03).

In the comparison of Remember vs. Imagine cues, the cluster-based test revealed a similar significant difference as in the comparison to Forget cues. Remember cues elicited greater amplitude than Imagine cues between 120 and 550 ms over central channels (*p* < 0.01). A second cluster was also found with a left frontal distribution between 780 and 1000 ms, with Imagine cues eliciting higher amplitudes than Remember cues (*p* = 0.03).

Lastly, in the comparison of Imagine vs. Forget cues, the cluster-based test revealed a similar significant difference as in the previous two comparisons. Forget cues elicited greater amplitude than Imagine cues between 110 and 570 ms over central channels (*p* < 0.01). Additionally, a second cluster revealed a significant difference between conditions, with Imagine cues eliciting greater amplitude than Forget cues over left frontal channels between 810 and 1000 ms (*p* = 0.02).

In summary, ERP amplitudes over central-posterior channels from roughly 150-550 ms differentiated cue conditions, with Remember cues eliciting greater amplitues than Forget cues, and Forget cues eliciting greater amplitudes than Imagine cues. Additionally, Imagine cues elicited a later positivity over frontal sites, from roughly 800-1000 ms, that was not observed following Remember or Forget cues.

### Time Frequency Results

Cluster-based permutation tests were utilized to compare oscillatory activity generated by the Remember, Forget, and Imagine cues. First, Forget cues elicited greater power than Remember cues over frontal sites from 450 to 1000 ms (*p* < 0.01). The cluster spanned roughly 5 to 15 Hz in frequency. Second, a similar significant cluster was found when comparing Forget cues and Imagine cues, though spanning a larger frequency range, 5 to 28 Hz (*p* < 0.01). The cluster spanned 500 to 1000 ms, and extended over both posterior and frontal channels, with Forget cues eliciting higher power than Imagine cues. Finally, when comparing Remember and Imagine cues, the cluster-based permutation test revealed a significant difference between conditions, with Remember cues eliciting greater power than Imagine cues (*p* < 0.01). The cluster extended 14-30 Hz, and spanned 500 to 900 ms with a posterior topography. These results suggest that the observed cluster in the Forget and Imagine comparison resulted from two effects: a power increase at lower frequencies elicited by Forget cues, and a power decrease at higher frequencies elicited by Imagine cues.

Figure 4 presents the time-frequency results, depicting the observed clusters. In summary, compared to Remember cues, Forget cues elicited *greater* frontal power at lower frequencies (in the theta / alpha range), whereas Imagine led to *reduced* power at higher frequencies (in the alpha / beta range) over posterior channels.

**Figure 4.**
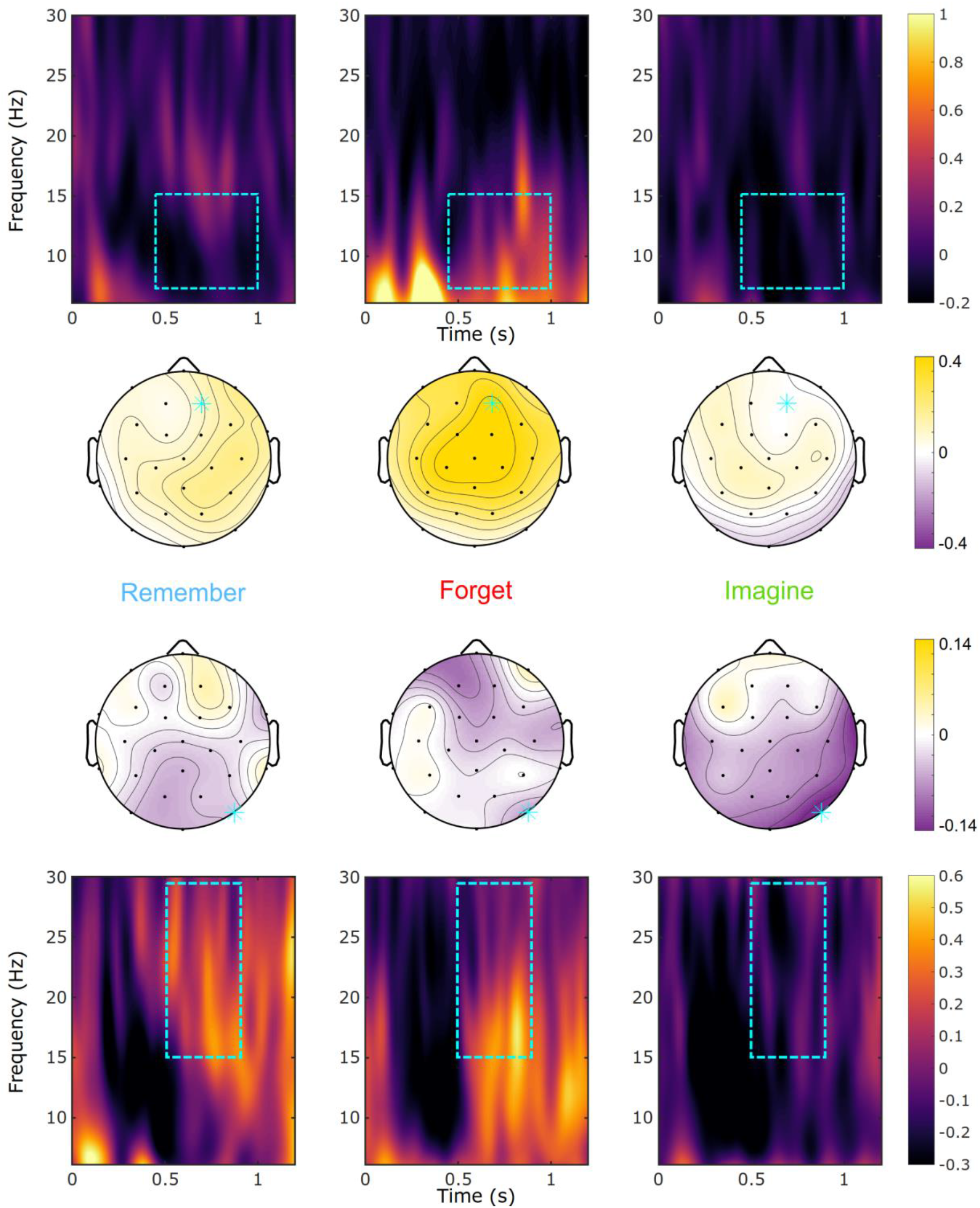
Time-frequency results. The top half shows highlights the observed frontal cluster for Forget cues, while the bottom half highlights the posterior cluster for Imagine cues. Topographic plots show the averaged time-frequency power in the corresponding box of the time-frequency plots, while time-frequency plots show power over time at the corresponding highlighted channel of the topography plots.

### RSA Results

RSA generalization matrices are presented in Figure 5. Cluster-based permutation tests on RSA generalization matrices revealed significant differences in item-cue similarity between conditions. First, a significant cluster was found in the comparison of Remember and Forget conditions (*p* = 0.02), and a similar cluster was found in the comparison of Imagine and Forget conditions (*p* < 0.01). Item-cue similarity was greater in the Remember and Imagine conditions compared to the Forget condition from roughly 100 to 200 ms post-item and 200 to 500 ms postcue. An additional cluster was found in the comparison of Imagine and Forget conditions (*p* < 0.01), with greater similarity in the Imagine condition roughly 400 to 600 ms post-item and postcue. A similar cluster was found in the comparison of Remember and Imagine conditions (*p* = 0.04), though in a slightly different time window (200-400 ms post-cue, 400-600 ms postencoding).

**Figure 5.**
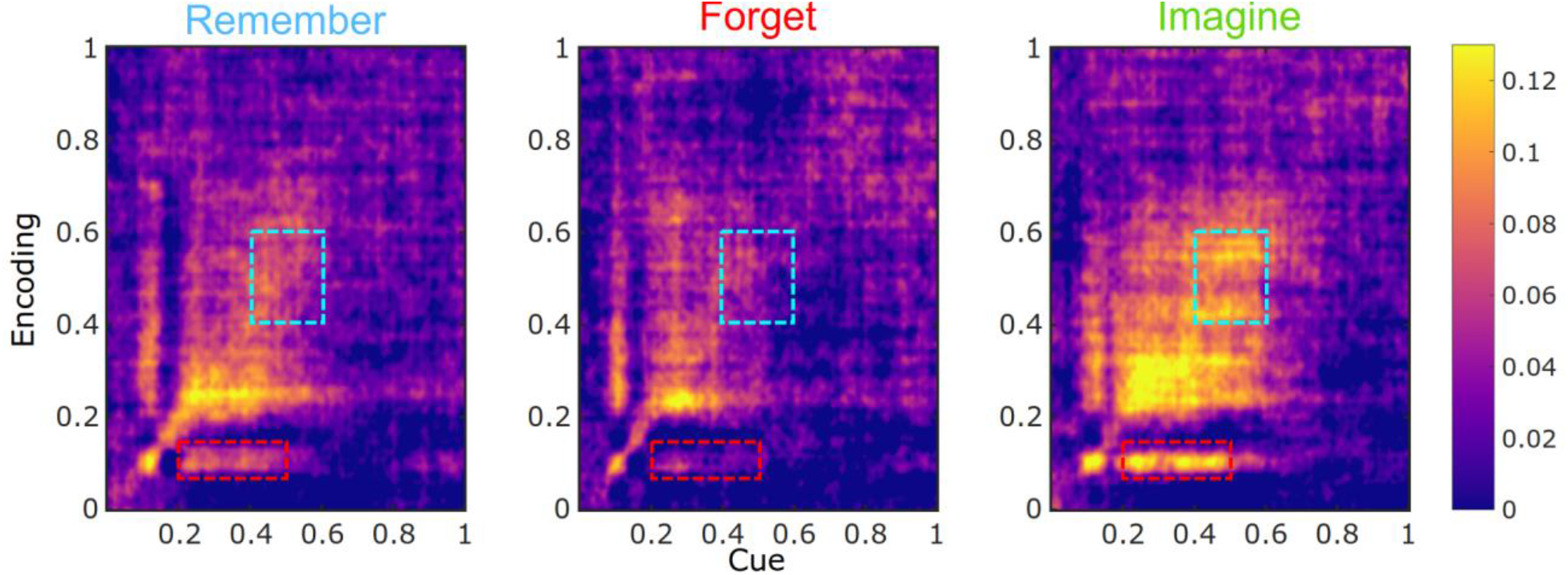
RSA results. Spatial generalization matrices for each condition are plotted. Time following the presentation of the item is plotted on the Y axis, time following presentation of the memory cue is on the X axis, and the color shows the magnitude of neural similarity. Identified clusters are highlighted in boxes, with the early cluster in red, and the later cluster in blue.

### Behavioral Prediction

The trial-level measurements of the previously identified neural clusters were submitted to a mixed-effect model predicting memory success (correct vs incorrect response). Specifically, mean signals were extracted from the previously identified significant clusters: ERP activity at central channels from 150-500 ms, ERP activity at left frontal channels from 800-1000 ms, time-frequency power (6-15 Hz) over frontal channels from 450-900 ms, posterior power (15-30 Hz) from 600-900 ms, average similarity from the early RSA cluster, and average similarity from the later RSA cluster. These 6 signals were entered as fixed effects, and the results of the analysis are reported in the following tables.

An initial model was run with cue (Remember, Forget, or Imagine) as a fixed effect, as well as the interaction with cue and each of the 6 brain signals. The fixed effects are summarized in Table 1. Significant fixed effects included the ERPs and frontal power signals, and interactions between cue and the ERP signals, as well as cue and late RSA.

**Table 1.** Predicting recognition accuracy from brain signals. Mixed effect model output is presented, with the fixed effects on the left in bold.

| Fixed Effect | Estimate | Std. Error | Z Value | *p* value |
| --- | --- | --- | --- | --- |
| **Cue** | 1.36 | 0.12 | 11.48 | **< 0.01** |
| **Central ERP** | 0.31 | 0.08 | 3.95 | **< 0.01** |
| **Frontal ERP** | 0.18 | 0.06 | 2.90 | **< 0.01** |
| **Frontal Power** | -0.20 | 0.07 | -2.90 | **< 0.01** |
| **Posterior Power** | 0.02 | 0.08 | 0.24 | 0.81 |
| **Early RSA** | -0.25 | 0.19 | -1.29 | 0.20 |
| **Late RSA** | 0.29 | 0.19 | 1.49 | 0.14 |
| **Cue\*CentERP** | 0.37 | 0.18 | 2.09 | **0.04** |
| **Cue\*FrntERP** | -0.47 | 0.19 | -2.56 | **0.01** |
| **Cue\*FrntPow** | 0.05 | 0.22 | 0.24 | 0.81 |
| **Cue\*PostPow** | 0.06 | 0.21 | 0.29 | 0.77 |
| **Cue\*Early RSA** | -0.77 | 0.54 | -1.42 | 0.16 |
| **Cue\*Late RSA** | -1.40 | 0.58 | -2.42 | **0.02** |

Given the significant interactions with cue, we ran mixed-effects models predicting accuracy for trials in each cue condition separately to better understand how these signals relate to successful remembering vs intentional forgetting. The fixed effects from these models are summarized in Table 2. For Remember cue items, only ERP activity was predictive of accuracy. Namely, higher ERP activity was associated with better memory accuracy. In contrast, for Forget cue trials, frontal power was predictive of accuracy. That is, higher frontal power was associated with lower accuracy, indicating that higher front power was more likely to contribute to successful intentional forgetting (e.g., low accuracy indicates successful DF in the Forget condition). Lastly, for Imagine cue trials, the effects of ERP and late similarity RSA were significant, both of which were positively associated with recognition accuracy.

**Table 2.**
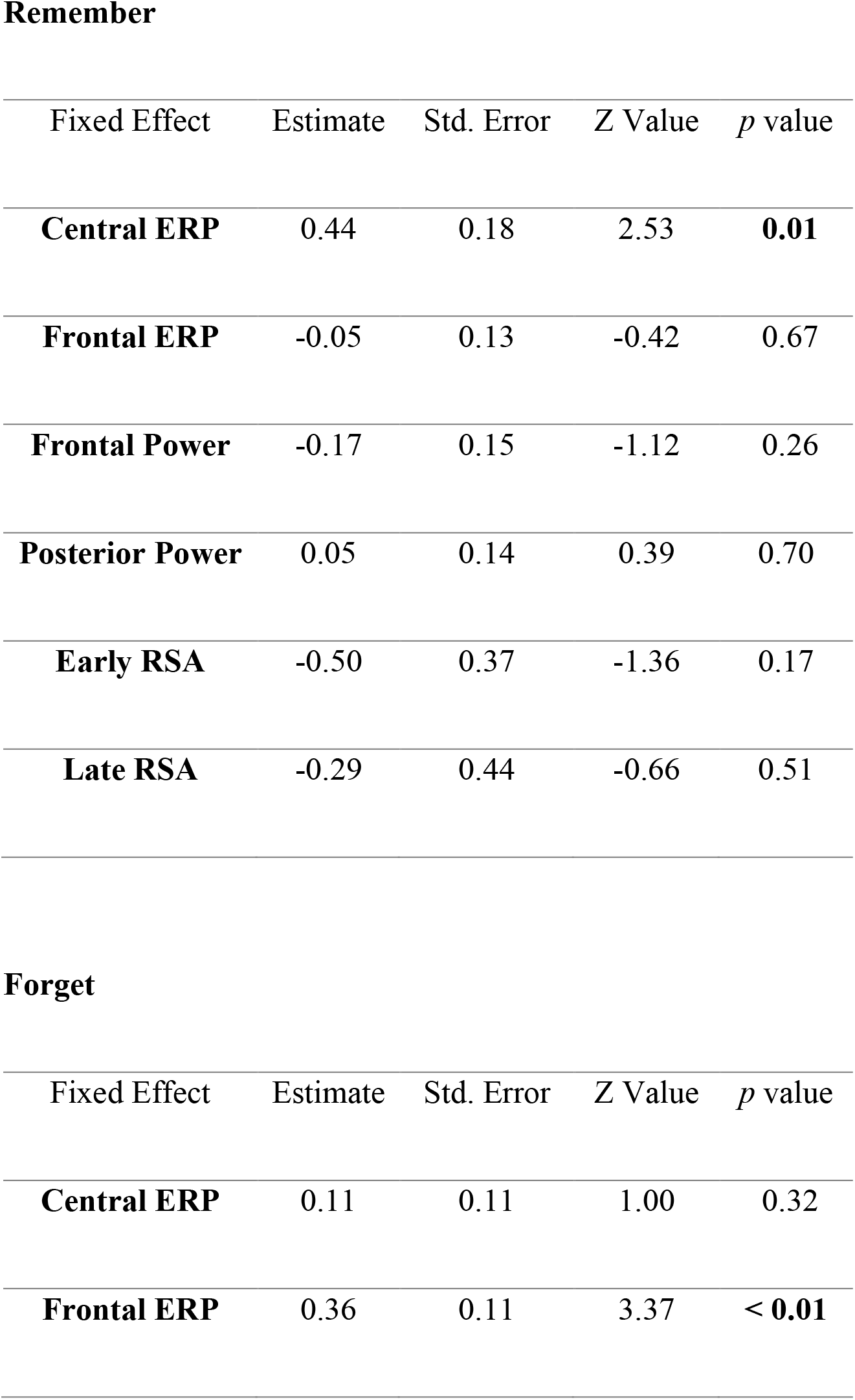
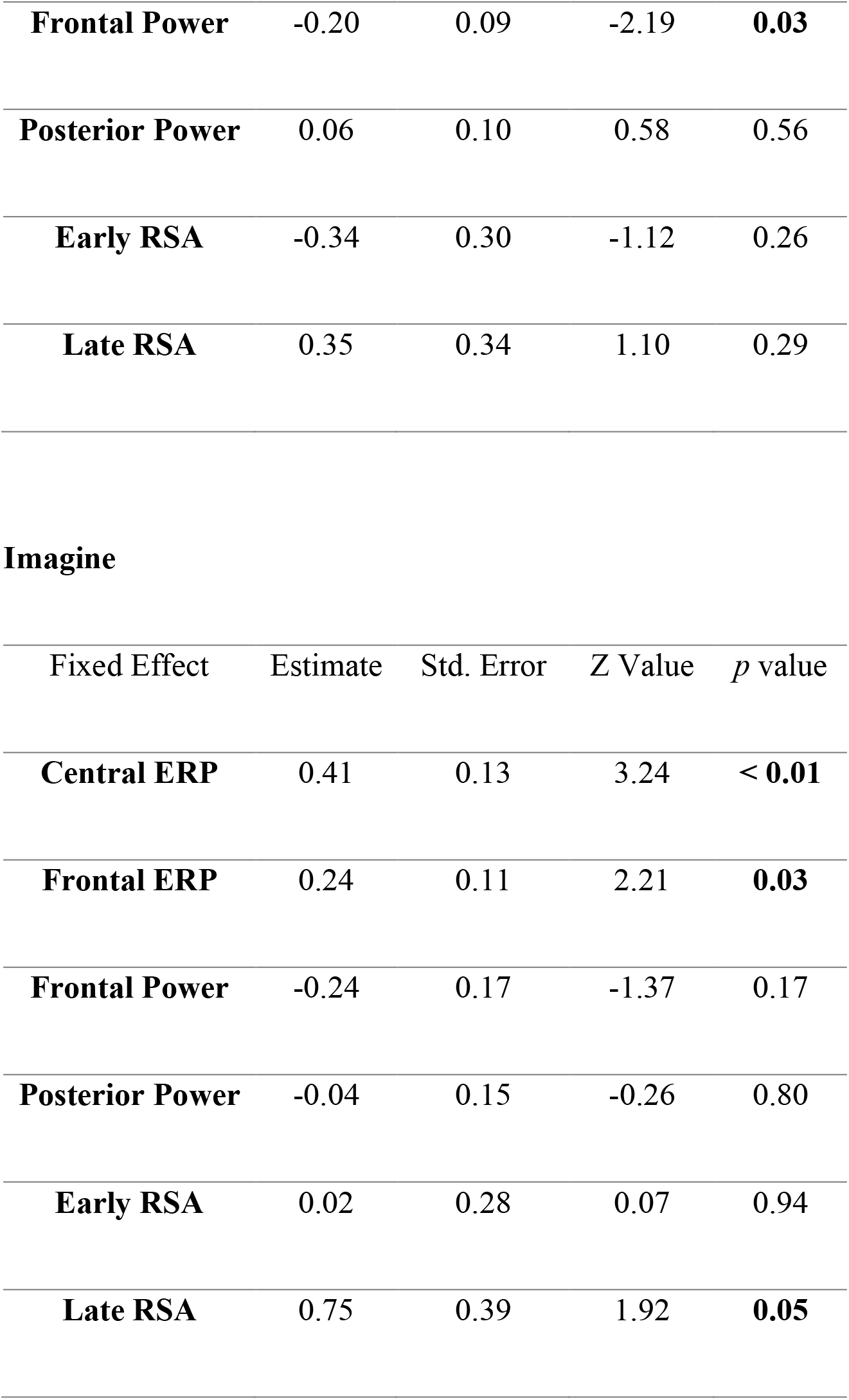
Predicting recognition accuracy from brain signals in each of the three cue conditions. Each table shows the regression results for a particular Cue Condition.

**Remember**
| Fixed Effect | Estimate | Std. Error | Z Value | *p* value |
| --- | --- | --- | --- | --- |
| **Central ERP** | 0.44 | 0.18 | 2.53 | **0.01** |
| **Frontal ERP** | -0.05 | 0.13 | -0.42 | 0.67 |
| **Frontal Power** | -0.17 | 0.15 | -1.12 | 0.26 |
| **Posterior Power** | 0.05 | 0.14 | 0.39 | 0.70 |
| **Early RSA** | -0.50 | 0.37 | -1.36 | 0.17 |
| **Late RSA** | -0.29 | 0.44 | -0.66 | 0.51 |

**Forget**
| Fixed Effect | Estimate | Std. Error | Z Value | *p* value |
| --- | --- | --- | --- | --- |
| **Central ERP** | 0.11 | 0.11 | 1.00 | 0.32 |
| **Frontal ERP** | 0.36 | 0.11 | 3.37 | **< 0.01** |
| **Frontal Power** | -0.20 | 0.09 | -2.19 | **0.03** |
| **Posterior Power** | 0.06 | 0.10 | 0.58 | 0.56 |
| **Early RSA** | -0.34 | 0.30 | -1.12 | 0.26 |
| **Late RSA** | 0.35 | 0.34 | 1.10 | 0.29 |

| Fixed Effect | Estimate | Std. Error | Z Value | *p* value |
| --- | --- | --- | --- | --- |
| **Central ERP** | 0.41 | 0.13 | 3.24 | **< 0.01** |
| **Frontal ERP** | 0.24 | 0.11 | 2.21 | **0.03** |
| **Frontal Power** | -0.24 | 0.17 | -1.37 | 0.17 |
| **Posterior Power** | -0.04 | 0.15 | -0.26 | 0.80 |
| **Early RSA** | 0.02 | 0.28 | 0.07 | 0.94 |
| **Late RSA** | 0.75 | 0.39 | 1.92 | **0.05** |

## Discussion

The aim of the current study was to investigate the mechanisms underlying two different strategies of intentional forgetting – namely, directed forgetting and thought substitution. Previous studies using list-method or think / no-think paradigms have shown that both strategies can lead to successful forgetting, and may recruit dissociable neural mechanisms, but no study to date has implemented an item-method paradigm with thought substitution cues to examine the utility of this strategy for forgetting individual items or episodes. We addressed this gap in the literature by employing a novel item-method paradigm with Remember, Forget, and Imagine cues while EEG was recorded. This permitted not only comparing forgetting rates of directed forgetting and thought substitution, but also investigating the neural mechanisms involved in these strategies. Using this approach, we were able to provide evidence that thought substitution is a viable strategy for intentional forgetting of individual items at a short time-scale. Namely, Forget and Imagine cues produced similar rates of forgetting, though Forget cues led to greater forgetting than Imagine cues. This differed from the aforementioned list-method studies, where comparable rates were found for directed forgetting and thought substitution (Sahakyan & Kelley, 2002; Sahakyan & Delaney, 2003). Thus, while both strategies can produce forgetting, directed forgetting may be more effective for individual items compared to thought substitution.

EEG was recorded during the experiment, allowing us to scrutinize the neural mechanisms underlying these strategies. Importantly, we employed predictive modeling to tie the observed differences between cue conditions in brain activity to trial-level recognition success. An early, sustained central ERP response differentiated cue conditions, and was predictive of accuracy for Remember cues. This finding of greater amplitude ERPs for Remember vs Forget cues is similar to other work examining ERPs to memory cues (Gallant & Dyson, 2016; Hsieh, Hung, Tzeng et al., 2009; Paz-Caballero, Menor & Jiménez, 2004; van Hooff & Ford, 2011). The posterior scalp distribution of the effect was similar to that of a P300 component (Polich & Kok, 1995), though with an earlier onset than is usually observed, potentially suggesting a P2 component difference as well (Luck & Hillyard, 1994). P2 amplitude modulations have been observed for repeated stimuli (Curran & Dien, 2003; Evans & Federmeier, 2007), while numerous studies have implicated P300 amplitude modulations as indexing goal-directed detection and subsequent memory processing of target stimuli (Polich, 2007). A difference in amplitude in these components that was predictive of accuracy for Remember cues suggests that Remember cues served as “targets” for participants; thus, participants may have devoted attentional resources and engaged in rehearsal processes when a Remember cue was presented, leading to a “repetition” of the stimulus and a P2 amplitude difference, as well as a P300 amplitude difference. Importantly, this result reveals that fairly rapid neural processes following remember cues contribute to successful memory encoding.

We also found a second ERP response that differentiated memory cues – a later frontal positivity that appeared more pronounced for Imagine cues than for Remember or Forget cues. Surprisingly, the amplitude of this ERP was predictive of memory accuracy for both Forget and Imagine cues, suggesting recruitment of a similar process for both of these conditions. Previous EEG studies have found enhanced frontal activity for Forget cues compared to Remember cues (Hauswald, Schulz, Iordanov et al., 2011; Paz-Caballero & Menor, 1999; Paz-Caballero, Menor & Jimenez, 2004; van Hooff & Ford, 2011), and fMRI data suggests the prefrontal cortex plays a role in intentional forgetting (Benoit & Anderson, 2012), although this relationship is less apparent for thought substitution. In the current study, greater amplitude of this ERP was positively associated with recognition accuracy, as opposed to negatively associated with accuracy (which would indicate successful intentional forgetting), a result at odds with the previous findings. One explanation is that this ERP does not index forgetting or inhibition mechanisms, but are related to some other process during encoding. For instance, other memory studies have identified left frontal neural activity in relation to semantic or associative encoding of information compared to more item-specific types of encoding (Fletcher, Shallice & Dolan, 2000; Gabrieli, Poldrack & Desmond, 1998; Köhler, Paus, Buckner et al, 2004). Thus, one possibility is that the left frontal ERP indexes selection and engagement of particular processing strategies, and participants engage in differential strategies following Forget or Imagine cues compared to Remember cues, leading to a difference between conditions in the current results.

Turning to the results of the time-frequency analysis, we also replicated prior results with our finding that frontal theta power was predictive of successful forgetting in response to Forget cues, with greater theta power related to more forgetting. In an intracranial EEG study of directed forgetting (Oehrn et al., 2018), Forget cues elicited greater theta power in the DLPFC, and this was related to successful forgetting. Additionally, the authors performed a granger causality analysis that showed significant information flow from the DLPFC to the hippocampus after Forget cue instructions. Given the pre-existing link between DLPFC and hippocampus in the theta band during memory formation (Benchenane, Peyrache, Khamassi et al., 2010; Gruber, Hsieh, Staresina et al., 2018), as well as memory retrieval (Anderson, Rajagovindan, Ghacibeh et al., 2010), it is possible that Forget cues recruit the same neural pathways, and engage frontal control mechanisms to inhibit or shut down memory encoding in the hippocampus. While EEG lacks the spatial resolution to identify specific neural generators of signals measured on the scalp, the frontal topography of our observed theta effect is suggestive of a DLPFC-generated signal, in line with the previously found result. Importantly, the topography and the direction of this effect differed compared to the previously described left frontal ERP effect, and the frontal theta effect was not a significant predictor of accuracy for the Imagine condition, and thus these are likely different neural signals. Additionally, no such frontal theta effects were found in response to Imagine cues, suggesting this inhibitory frontal control response is not engaged during thought substitution.

In contrast to Forget and Remember cues, Imagine cues led to a decrease in alpha power, a similar result to what was found in a list-method study (Pastötter, Bäuml & Hanslmayr, 2008). Interestingly, this alpha decrease was not predictive of successful forgetting, and thus may have been linked to a more general mechanism of thought substitution. Alpha band decreases have been reported in other memory studies, and have been related to successful memory encoding (Hanslmayr, Spitzer & Bäuml, 2009; Hanslmayr, Staudigl & Fellner, 2012; Sederberg, Kahana, Howard et al., 2003), which may be related to the richness or number of details encoded about a memory trace, though these conclusions were based on standard Dm analyses as opposed to predictive modeling. However, it is unclear why this would on be observed in response to Imagine cues, and not Remember cues as well, where the goal is to encode the stimulus. An alternative explanation is that Imagine cues led to focused attention toward the substituting thought and away from the stimulus, leading to changes in alpha power. Alpha power has been shown to increase during periods of introspection and mind wandering (Arnau, Löffler, Rummel et al., 2020; Boudewyn & Carter, 2018; Compton, Gearinger & Wild, 2019), which may seem similar to thought substitution (e.g., Delaney et al., 2010); however, mind wandering reflects off-task thought or a lapse in sustained attention, whereas thought substitution may in fact require focused attention away from the processing of the stimulus. This attention toward the substituting thought may always be engaged during thought substitution, but is not the mechanism most related to successful forgetting of the stimulus.

For the third and final analysis, we employed representational similarity analysis to examine the similarity of neural activity during processing of the item and neural activity involved in processing the cue. This strategy has been used in one other study to examine active inhibition following Forget cues (Fellner, Waldhauser & Axmacher, 2020). Interestingly, we did not replicate this study’s results, which may have been due to differences in stimulus materials or timing parameters of item and cue presentation between the two experiments. However, we found a cluster of cue-item similarity that was predictive of successful forgetting following Imagine cues, with less similarity related to greater forgetting. One explanation is that the emergence of the neural representation of the item during the cue period reflected intrusion of item processing or rehearsal, i.e. failure of thought substitution. However, by this explanation, the cue-item similarity would likely also be predictive of success following Forget cues, as some item processing could occur when intentional forgetting failed. Alternatively, this relationship between similarity and recognition memory performance following Imagine cues may indicate a shift in context produced by the substituting thought. Previous list-method studies suggest that thought substitution causes a change in mental context, leading to difficulty successfully retrieving the items from the list (Sahakyan & Kelley, 2002). Here, if thought substitution successfully shifted context, then the pattern of neural activity during the cue period would differ more from the item period activity, as the contexts would not overlap as much. This would produce difficulty in retrieving the item later, as the context associated with the item would differ from the context of the encoding period, and thus reinstating the encoding context would not serve as a useful cue for recognition of the item. Thus, while directed forgetting may act through direct suppression or top-down control of item processing, thought substitution may act through shifting context, causing poor item-to-context binding.

In sum, our results demonstrate that participants recruit different strategies, as well as neural mechanisms, to perform intentional forgetting or thought substitution. Additionally, our neurobiological results in conjunction with the behavioral result of greater forgetting for Forget cues compared to Imagine cues also provides evidence against a classic theoretical account of the mechanism of item-method directed forgetting. Namely, the traditional interpretation of the item-method DF effect, known as the *selective rehearsal* account, emphasizes passive processes that involve removing the Forget items from rehearsal in working memory, while Remember items remain and are more elaborately encoded (e.g., Bjork, 1970; 1972; MacLeod, 1975; Basden, Basden, & Gargano, 1993). This explanation focuses on processes acting on Remember items; namely, the Remember cues lead to additional processing and encoding items, while Forget cues simply do not engage these processes. Contrary to this view, other research suggests that more active processes such as inhibition may underlie intentional forgetting (Anderson & Hanslmayr, 2014). Some work has supported this *inhibition account*; for instance, studies have shown slower reaction times to a secondary task performed during execution of the Forget instruction compared to the Remember instruction, as well as larger inhibition of return following the Forget cue, indicating that forgetting is effortful (e.g., Fawcett & Taylor, 2008; 2012; Taylor, 2005). Recent eyetracking work has also supported an active inhibitory account of directed forgetting (Whitlock, Lo, Chiu et al., 2020). In the current study, participants were unlikely to engage in rehearsal during Imagine cues, as they were thinking of the given cue, and recognition performance in the Imagine condition was indeed lower than the Remember condition. If intentional forgetting elicited by Forget cues also simply reflected a lack of rehearsal, then equivalent forgetting as the Imagine condition would be expected. However, Forget cues led to even greater forgetting than Imagine cues, suggesting Forget cues are not simply the absence of rehearsal, but instead engage an active inhibition mechanism. Additionally, only Forget cues elicited frontal theta activity that was predictive of forgetting, supporting this active inhibition account.

Our results provide novel evidence that thought substitution can be employed in an item-method design, but recruits different neural mechanisms compared to intentional forgetting. This opens the door for future studies to better characterize the differences between these two strategies, both across different populations that may differ in their successful usage of these strategies, as well as across the conditions or parameters that modulate successful intentional forgetting.

## Acknowledgements

This work was supported by a Beckman Postdoctoral Fellowship granted to R.J.H.

## Competing Interests

The authors declare no competing financial or non-financial interests in this work.

## Notes

### Competing Interest Statement

The authors have declared no competing interest.

